# Locomotor compromises maintain group cohesion in baboon troops on the move

**DOI:** 10.1101/2020.10.15.340711

**Authors:** Roi Harel, J. Carter Loftus, Margaret C. Crofoot

## Abstract

When members of a group differ in locomotor capacity, coordinating collective movement poses a challenge: some individuals may have to move faster (or slower) than their preferred speed to remain together. Such compromises have energetic repercussions yet research in collective behavior has largely neglected locomotor consensus costs. Here we integrate high-resolution tracking of wild baboon locomotion and movement with simulations to demonstrate that size-based variation in locomotor capacity poses an obstacle to collective movement. While all baboons modulate their gait and move-pause dynamics during collective movement, the costs of maintaining cohesion are disproportionately borne by smaller group members. Although consensus costs are not distributed equally, all group-mates do make locomotor compromises, suggesting a shared decision-making process drives the pace of collective movement in this highly despotic species. These results highlight the importance of considering how social dynamics and locomotor capacity interact to shape the movement ecology of group-living species.

## Introduction

Group-living animals incur consensus costs when they compromise their own preferred course of action to remain in contact with other members of their group (Conradt & Roper 2005; Pyritz *et al*. 2010). When group members vary in their physical characteristics (e.g., body size), consensus costs may be particularly high, as physiological differences can introduce significant conflicts of interest among group-mates. Differences in locomotor capacity – the ability of an organism to move through its environment – may pose particularly severe challenges to behavioral coordination in heterogeneous groups. Locomotor capacity, which is dependent on a range of morphological features including body weight and limb length, affects the energetic costs of movement and therefore serves as a major driver of movement decisions (Perrigo 1987; Lees *et al*. 2012; Halsey 2016). Studies of several species of terrestrial animals revealed that individuals have a preferred travel speed (Pennycuick 1975) which is hypothesized to maximize energy efficiency (Hoyt & Taylor 1981).

Because physical characteristics, such as limb length and body mass, shape preferred travel speeds (Heglund & Taylor 1988), variation among individuals in body size will lead to differences in optimal stride frequencies and travel speeds within groups. How do groups maintain cohesion during collective movement when faced with such inter-individual differences?

The locomotor choices that individuals make with respect to stride frequency and length have important effects on their energetic cost of transport (bipeds: Muro-de-la-herran *et al*. 2014; Maculewicz *et al*. 2016; quadrupeds: Heglund *et al*. 1982; Dewhirst *et al*. 2017). Despite the obvious potential for differences in preferred travel speed and stride frequency to introduce behavioral and energetic conflicts of interest when individuals move together as a group, our understanding of the impact of locomotor capacity on collective movement is limited (Jolles *et al*. 2020). In social species, high inter-individual variation in locomotor capacity is expected to impose significant costs if group members must alter their patterns of movement in order to maintain group cohesion (Delgado *et al*. 2018; Sankey *et al*. 2019). Cohesion could be maintained in two ways: individuals with higher locomotor capacity can slow down or pause to allow other group members to catch up, or individuals with lower locomotor capacity can travel faster or pause less frequently to keep up with their group-mates. In either case, some group members pay a cost. Faster animals who slow down or pause and wait pay an opportunity cost because they commit additional time to transit that could have otherwise been devoted to other activities such as feeding. Individuals who speed up to remain with their group, or who take fewer breaks during travel, increase their energetic cost of locomotion. On the other hand, if group members fail to coordinate, the resulting increase in group spread is expected to be costly, with individuals potentially experiencing greater exposure to predators and suffering from reduced information transfer (Lindström 1999; Ronget *et al*. 2018).

To test how members of heterogeneous groups maintain cohesion during collective movement, we fit GPS collars with integrated tri-axial accelerometers to the majority of members of a wild olive baboon (*Papio anubis*) group. Olive baboons live in cohesive groups of up to 150 individuals that travel together throughout the day in search of resources. Because they are sexually dimorphic and maintain stable mixed-age groups, they exhibit large within-troop variation in body size (Ray & Sapolsky 1992; Dunbar 2013).

We first tested whether differences in body size translated into differences in stride frequency, as well as vectorial dynamic body acceleration, an established proxy for energetic expenditure (Quasem *et al*. 2012, Wilson *et al*. 2020). We then investigated how fine-scale movements preserved group cohesion, and, in doing so, identified decision rules that might generate the observed patterns. Baboons group-mates do not benefit equally from their membership in their troop (Barton & Whiten 1993; Silk *et al*. 2009), and thus we hypothesized that individuals who had more to gain from group membership would be willing to incur additional locomotor costs to keep the group together. Specifically, due to their greater vulnerability to predators (Cowlishaw 1994), we predicted that smaller baboons would be more sensitive to their spatial positioning, make larger behavioral compromises, and bear more of the costs of maintaining group cohesion, compared to larger group members.

## Methods

### Data collection

Simultaneous tracking data were collected from 25 wild olive baboons (*Papio anubis*) belonging to a single group at the Mpala Research Centre in Laikipia, Kenya (Figure 1). The GPS data have formed the basis for several studies on individual positioning (Farine *et al*. 2017), collective movement (Farine *et al*. 2016; Strandburg-Peshkin *et al*. 2017) and consensus decision-making (Strandburg-Peshkin *et al*. 2015). Collar units recorded location estimates continuously at a 1 Hz sampling interval and tri-axial acceleration data at 12 Hz during daylight hours (06-18h) from August 1^st^ to September 2^nd^, 2012. While individuals were chemically immobilized and being fit with telemetry collars (see Strandburg-Peshkin *et al*. 2015 for details on capture methodology), the length of each individual’s front leg was measured (dorsal most point of the scapula to the carpus; hereafter referred to as “leg length”). Collared individuals consisted of 80% (23/29) of the adult (N = 13) and subadult (N = 10) members of the group, as well as two juveniles. Two of the adult individuals were removed from the analyses due to missing body measurements or inconsistencies in acceleration data. Focal video recordings of the behavior of collared individuals were collected and coded to identify periods of stationary and non-stationary behavior. In total, 20 minutes of non-stationary behavior and 180 minutes of stationary behavior was recorded in the field.

**Figure 1.**
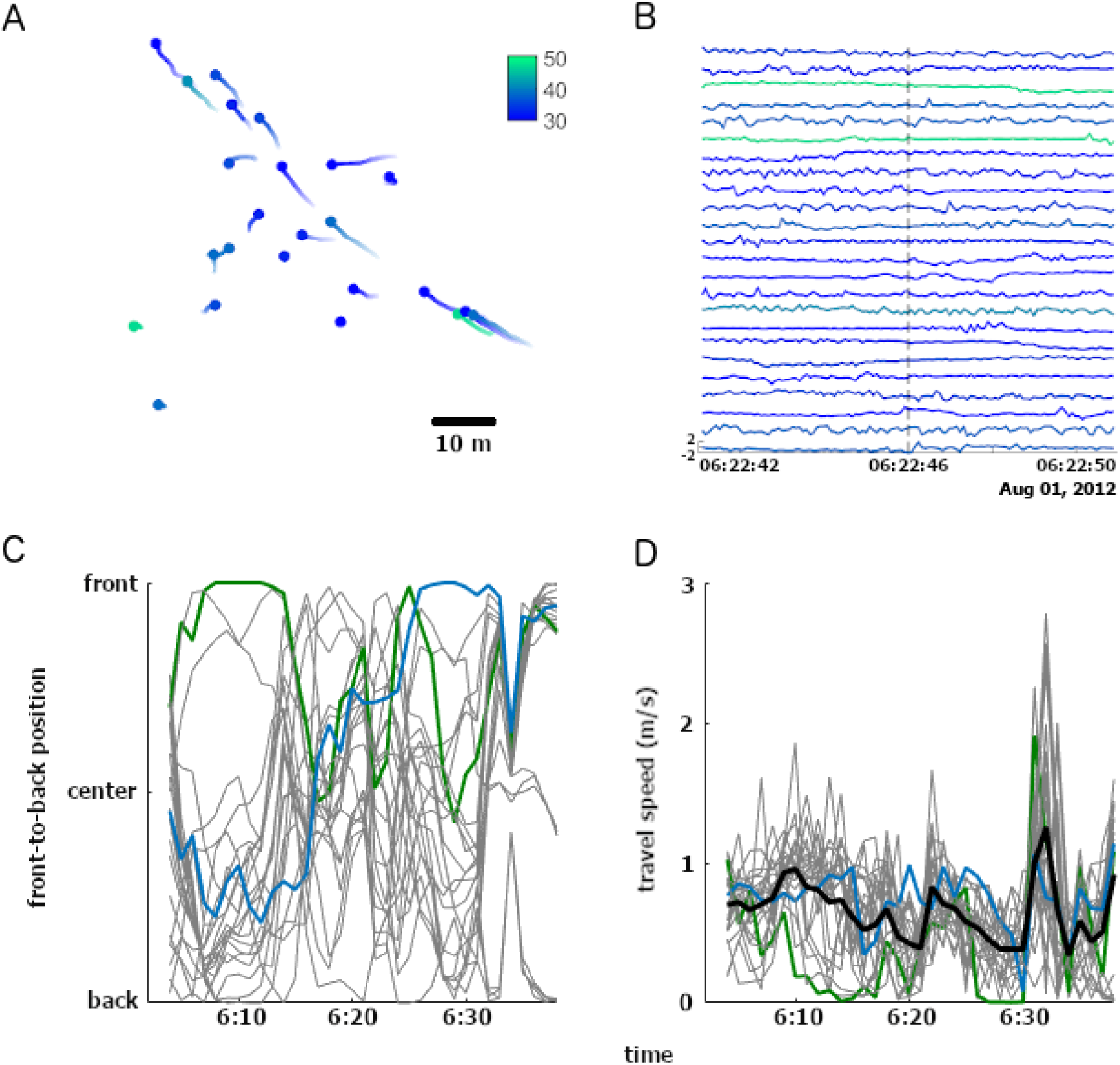
Visualizing locomotor parameters of baboons during a segment of movement. A snapshot of (a) the locations of baboons at time *t*, represented by circles, with tails stretching back to individuals’ locations at *t*-10s, and (b) the heave axis acceleration - with peaks representing footfalls – of all individuals, show variation in baboons’ move and pause activity states, as well as in their stride frequencies. Over a 40-minute period, (c) individuals’ position within the group, relative to the direction of group movement, as well as (d) individuals’ speeds during group travel, are highly variable. In (a) and (b) individuals with longer leg length than average are represented in green, and shorter than average, in blue. The blue and green colored lines on (c) and (d) highlight the patterns of two individuals. The black line in (d) represents the travel speed of the group centroid.

### Daily travel distance and displacement

To assess how movement patterns, vary with body size, we calculated the daily travel distance and maximum displacement from the sleeping site of each group member, as well as of the group’s centroid. Daily travel distance is a widely used measure of animal movement but is strongly affected by sampling frequency (Rowcliffe *et al*. 2012). For this reason, and to avoid the accumulation of GPS positional error inflating our estimates, we calculated daily travel distances after discretizing the data to 5-meter resolution (Strandburg-Peshkin *et al*. 2017). Maximum displacement from the sleeping site was measured as a straight-line distance between the group’s morning sleeping site and the most distant position visited on that day. We used linear mixed models (LMMs) to estimate the effects of leg length on (1) daily travel distance and (2) daily displacement, taking into account individual identity as a random effect and temporal autocorrelation between days using an autoregressive (AR1) component in both of the models (Pinheiro & Bates 2000).

### Group movement parameters

Group activity state was classified into two categories, stationary and non-stationary, based on changes in the displacement of the group centroid. Group travel bouts were classified using a change point detection algorithm (Lavielle 2005) on the centroid displacement speed. The speed and heading of the group’s centroid were calculated at 10 second intervals. Individuals’ relative positions within the group on the front-to-back axis were determined by multiplying their x-y locations by a rotation matrix based on the heading of the centroid. The resulting values were rescaled such that – regardless of group spread – 1 represents being at the front, 0 represents the center, and −1 represents the back of the group (Figure 1C). For some analyses, rather than front-to-back scaled values, individuals’ positional rank relative to group mates were used.

### Individual behavior and movement parameters

Individual’s activity state was inferred with a support vector machine following Fehlmann *et al*. 2017. Acceleration and location variables were time-matched with videos to obtain a labelled dataset. The most important features to classify the two activity states were heave peak frequency, and heave amplitude and the heave maximum power spectral density. We applied a Hampel filter to the acceleration data to remove spikes (Dewhirst *et al*. 2017) that were likely caused by direct physical strikes to the collar unit and trained the algorithm using ground-truthed labels derived from the video recordings. The algorithm distinguished moving and non-moving activity states, exhibiting an accuracy of 0.92.

We then estimated stride frequency based on the timing of heave-axis (i.e. baboons’ dorsal-ventral axis) peaks (Dewhirst *et al*. 2017) (Figure 1B). For each individual, we estimated a characteristic stride frequency by measuring her/his average stride frequency while moving alone, i.e. during periods when the individual was moving, but the rest of its group-mates were stationary, presumably engaged in foraging, socializing, or resting. Thus, the characteristic stride frequency of each individual represents the average stride frequency which she/he chooses, independent of social influences. We then tested for a correlation between individuals’ characteristic stride frequencies and their leg lengths during both single-individual and group movement.

The vectorial dynamic body acceleration (VeDBA) of each group member was calculated using data from tri-axial accelerometers, following Halsey *et al*. 2009 and Wilson *et al*. 2020. Derivatives of dynamic body acceleration, such as VeDBA and ODBA (overall dynamic body acceleration), are proxy measures for movement-based energetic expenditure that has been validated for several quadrupedal taxa (Halsey *et al*. 2009; Qasem *et al*. 2012; Williams *et al*. 2015; Wilson *et al*. 2020). We used a LMM to compare individuals’ VeDBA values averaged over 10 second intervals during single-individual and group movement.

### Modeling group spread and size-based segregation

To assess how the simple decisions that individuals make with respect to modulating their travel speed change the collective properties of their group, we compared model simulations to our observed data. We modelled three alternative scenarios for a group moving in a single dimension (Video 1) and compared the results of these simulations to the observed patterns of group spread, measured as the total Euclidean distance between the front-most and back-most group member. In the modelled scenarios, individuals (1) moved at their preferred speed (parameters included a characteristic speed for each agent drawn from the empirical data), (2) modulated their speed as function of their position in the group (parameters included a characteristic speed for each agent that varied as a function of their location in the group) or (3) moved at their preferred speed when group spread was low and modulated their speed as function of their position in the group when the spread exceeded a threshold value (parameters included a characteristic speed for each agent that varied as a function of their location in the group, and group spread). The durations of simulations were drawn from the distribution of the observed group travel bouts. We fit the parameters of speed modulation and the group spread threshold to find the values that best predict the observed data. Using these parameters, we then fit the models to the observed data and obtained an AIC (Akaike’s Information Criterion) value for each model.

The effect of body size variation on the emergence of spatial segregation was assessed by sampling the relative location of large and small individuals and calculating the front-to-back positional rank difference between the two size categories (divided by the group mean leg length). We used a linear mixed model (LMM) to test the effect of body size on the positional rank difference with the group movement event as a random effect, for both simulated and observed movement tracks.

### Individual behavior during group movement events

We examined how individuals of different sizes adjust their locomotor behavior in the context of group movement, as well as the energetic consequences of these fine-scale movement decisions. We limited this analysis to group travel bouts when the group centroid was moving for at least two minutes and data were available for at least 15 baboons, ensuring a reliable representation of the group when estimating its activity state and the front-to-back rank position of group-members.

Specifically, we assessed how a focal individual’s leg length, its position within the group (as a linear and quadratic term), the difference in leg length between the focal individual and its nearest neighbor (for a subset of cases in which the nearest neighbour distance was under 5-m), group speed, and group spread affected focal individuals’ (1) deviation in stride frequency from the characteristic stride frequency, and (2) VeDBA, measured in m/s^2. To account for the dynamic nature of both predictor and response variables within a travel bout, all measures were aggregated over 10 second intervals and each interval represented a single observation. The candidate generalized linear mixed models (GLMMs) included the above factors as main effects and also an interaction term between leg length and position in the group. All models accounted for individual identity and the cohesive group movement event as crossed random factors, and considered temporal autocorrelation by using an autoregressive moving average (ARMA) component. models were ranked according to AIC, a relative measure of parsimony, i.e. the balance between number of parameters and the fit of the model (Burnham & Anderson 1998).

We also examined how an individual’s position in the group affected the variation in its activity states. We calculated individuals’ ratio of time spent moving to time spent stationary, henceforth the “move:pause ratio.” We used GLMMs, with the move:pause ratio as the response variable, and focal’s leg length and its positions within the group as fixed effects. These models also accounted for individual identity and the group movement event as crossed random factors. Statistical analyses were performed in Matlab and R (R core team 2012) using the packages nlme and MuMin (Bartoń 2018; Pinheiro *et al*. 2018).

## Results

### Locomotor capacity varies with body size

Baboons in the study group varied substantially in size (leg length: mean = 38 cm, range = 31 to 51 cm), stride frequencies, daily movement patterns, and energetic costs. Individuals displayed characteristic stride frequencies that varied as a function of body size (Figure 2A): when the group as a whole was stationary and individuals moved independently, stride frequency was negatively correlated with leg length (Pearson correlation, r = −0.53, P = 0.01). Preferences for particular stride frequencies extended to the context of collective movement. When the group was moving cohesively (N = 96 travel bouts lasting 26 ± 2 minutes; mean + SE), larger individuals exhibited lower stride frequencies than smaller group-members (LMM, *b* ± SE = 0.013 ± 0.002 Hz, Wald *t* = 2.1).

**Figure 2.**
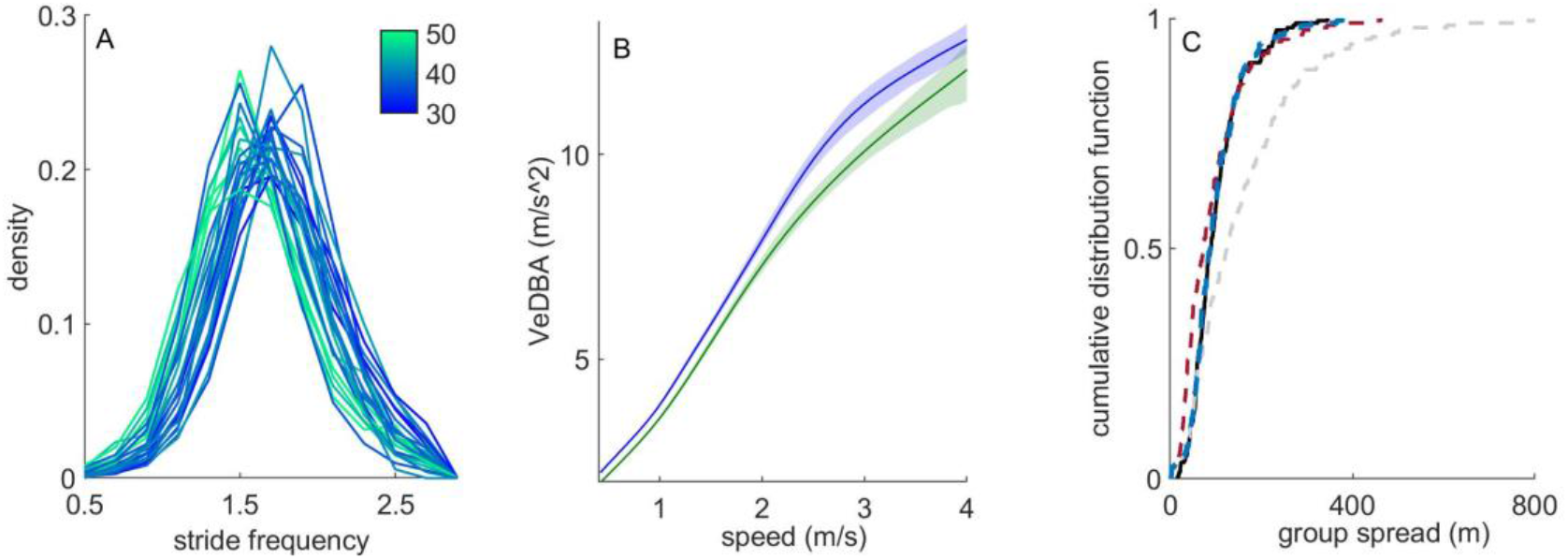
Body size affects stride frequency, energetic expenditure, and locomotor modulation. (a) Variation in individuals’ stride frequency while moving alone (i.e. during periods when the individual was moving, but the rest of its group-mates were stationary). Line color indicates leg length (in centimeters). (b) Differences in movement costs as estimated by VeDBA resulted in higher costs of movement for smaller individuals compared to large individuals travelling at the same speed. Individuals with longer leg length than average are represented in green, and shorter than average, in blue. (c) Cumulative distribution function for group spread under four alternative scenarios: observed (black solid), moving profile (gray dashed), position-dependent speed (green dashed), and position- and spread-dependent speed (blue dashed).

Baboons of different sizes also varied in their movement patterns; high-resolution GPS tracking revealed significant inter-individual variation in total daily distances travelled. The group as a whole – measured from the position of its centroid – traveled a mean of 10.2 (±2.7 SD) km per day, with most of that distance covered during long travel bouts, punctuated by periods when the group remained relatively stationary. Individual baboons travelled for 142±25 minutes each day, during which they covered 12.1±1.4 (mean±SD) km. Individual’s daily travel distance was negatively related to body size (Wald *t* = 1.91), and decreased 30 (±10 SD) m with each 1 cm increase in leg length. Only minor differences (±1%) were found between individuals’ daily maximum displacement from the sleeping site and that of the group’s centroid, reflecting their shared route.

Differences in individual locomotion and movement patterns had energetic consequences that disproportionately impacted smaller individuals, particularly when the group engaged in collective movement. Overall, VeDBA decreased with increasing body size (*b* ± SE = −0.13 ± 0.04 m/s^2 for each 1 cm change in leg length; Wald *t* = 2.68) and increased with travel speed (*b* ± SE = 6.15 ± 0.01, Wald *t* = 460.35, Figure 2B). Increases in travel speed had a larger impact on VeDBA of smaller baboons compared to their larger group-mates (*b* ± SE = 0.14 ± 0.03, Wald *t* = 3.89). VeDBA values were higher when individuals moved together compared to when they were moving and the group was stationary (*b* ± SE = 0.05 ± 0.005, Wald *t* = 9.74).

### Socially-mediated movement decisions maintain group cohesion

We compared observed patterns of group spread and size-based spatial segregation to patterns predicted by agent-based models where individuals varied their stride frequency as a function of their leg length. Models in which individuals moved without any modulation to their characteristic stride frequency overestimated group spread by up to 800% [ΔAIC = 260]. In contrast, incorporation of simple socially-based decision rules improved model performance. The addition of a single rule in which individuals vary their speed as a function of their position within the group provided a good fit to our observed data for long travel bouts, but underestimated group spread for short travel bouts [ΔAIC = 220]. The best fitting model incorporated position-dependent modulation of speed when group spread was larger than a threshold value (estimated to be 80 meters), but allowed individuals to move at their characteristic stride frequency when the group was highly cohesive (Figure 2C). These patterns align with our empirical data, where the mean deviation of group members from their characteristic stride frequency increased by a mean of 0.7 (±0.2 SD) % with 1 m increase in group spread (Wald *t* = 4.51; Figure 3E). Our model predicts that size-based segregation will emerge if group members move at their characteristic stride frequency and do not modulate their stride frequency based on their position in the group; in simulated travel bouts, front-to-back positional rank was positively associated with body size (LM; *b* ± SE = 2.71 ± 0.90, Wald *t* = 2.47). In contrast, we found no evidence for size-based segregation in our empirical travel bouts; the mean rank difference between large and small individuals (mean = 0.46, SD = 0.85) was not distinguishable from zero.

**Figure 3.**
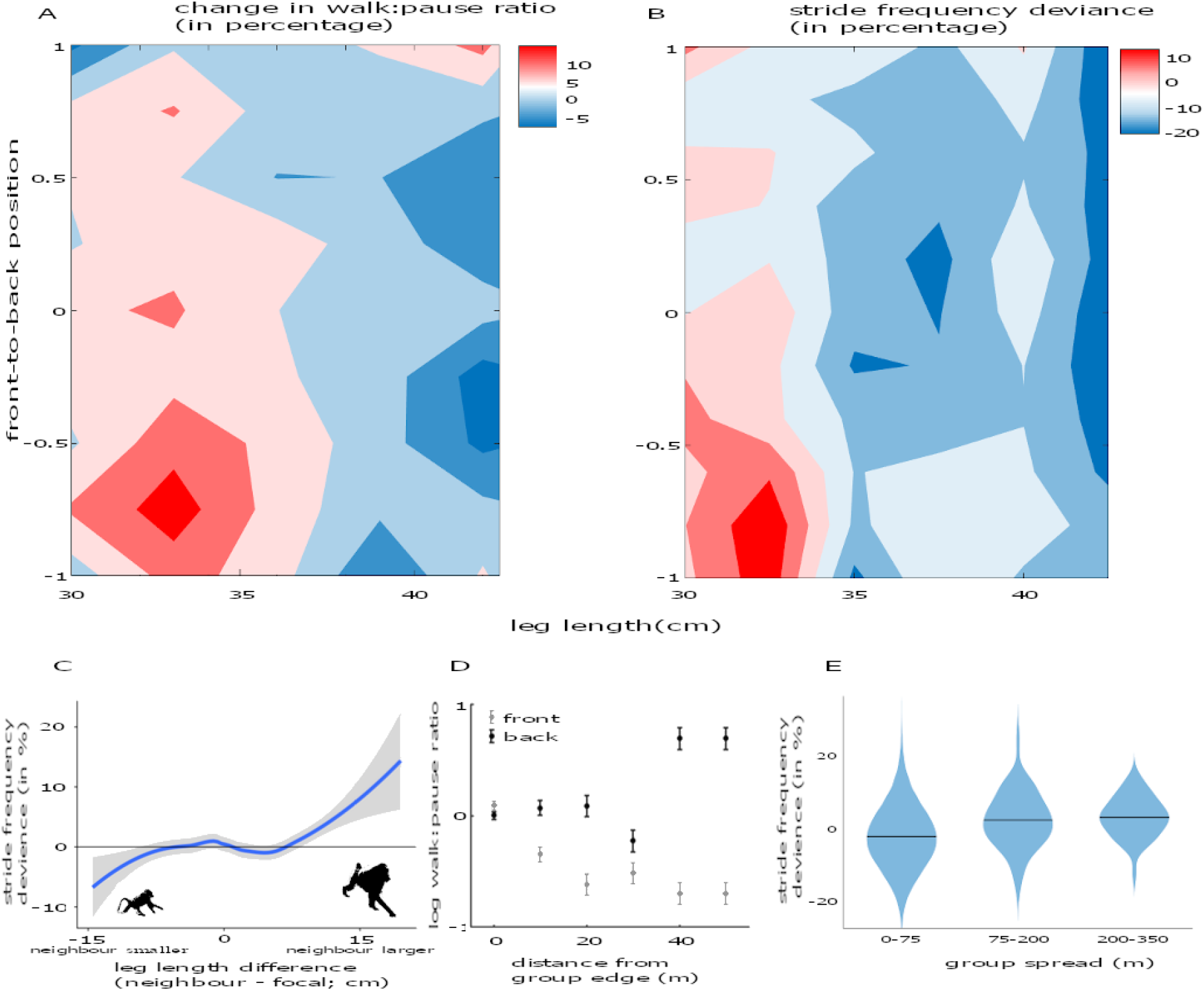
Context-dependent decision rules support the emergence of cohesion. (a) The move:pause ratio and (b) deviation of individuals’ stride frequencies from their characteristic stride frequencies varied depending on an individual’s leg length and front-to-back position in the group. (c) Differences in leg length between dyad members moving in proximity affects individuals’ behavior. In proximity of a larger neighbor, the focal individual increased its stride frequency, whereas, in proximity of smaller neighbor the focal individual decreased its stride frequency, but to a lesser degree. (d) The decision to change activity state was dependent on the position in the group as apparent from the log of the move:pause ratio. In the front, individuals were more sensitive to being separated from the group, and were more likely to change their behavior at a shorter distance from the rest of the group, than when individuals were at the back and separated from the rest of the group. (e) The deviation from characteristic stride frequency was lower when group spread was small (0-75 m) compared to larger group spread (75-150 m, 150-350 m) based on the trimodal distribution of group spread (Figure S1).

### Local decision rules support the emergence of cohesion

Baboons modulate their travel speed by varying their stride frequency and their move:pause ratios. An individual’s decision to adjust these fine-scale movement characteristics was sensitive to its social context. Individuals adjusted their stride frequency depending on the relative size of their nearest neighbor. Relative to their characteristic stride frequency, baboons increased their stride frequency when traveling in proximity (< 5 meters) to larger individuals, and decreased their stride frequency when in proximity to smaller individuals. However, the size of these behavioral adjustments was not equal; smaller individuals increased their stride frequency more than their larger neighbors decreased their stride frequency (LMM; *b* ± SE = 0.13 ± 0.04 %, Wald *t* = 2.68; Figure 3A).

Position within the group also influenced baboons’ movement decisions. While individuals in the front of the group maintained their characteristic stride frequency, baboons, regardless of size, increased their stride frequency when they were at the back (*b* ± SE = 1.6 ± 0.3 %, Wald *t* = 2.70; Figure 3B). However, the behavioral strategies of small and large individuals differed at the center of the group. In these central positions, smaller individuals increased their stride frequencies but larger individuals did not deviate from their characteristic stride frequency (significant interaction between the quadratic term of group position and leg length; 2 ± 0.2 %, Wald *t* = 2.14). Baboons changed their position within the group regularly (Figure 1C), maintaining the same positional rank along the front-back axis for an average of only 54.0 ± 14.1 seconds. Overall, smaller individuals exhibited higher move:pause ratios than their larger group-members (binomial GLMM, *b* ± SE = 0.46 ± 0.10, Wald *z* = 4.41), but this was especially true when they were at the back of the group (Figure 3A). All group members were less likely to move when they were at the front of the group, and more likely to move when they were at the back.

The spatial scale at which separation from the group prompted a change in an individual’s move:pause ratio differed depending on whether an animal had outstripped or had fallen behind the rest of the group. At the front of the group, the increase in move:pause ratios occurred when individuals got 20 m ahead of their group-mates, whereas individuals had to fall at least 40 m behind the rest of their group before increasing their move:pause ratios (Figure 3D).

## Discussion

In social species, variation in individual locomotor capacity complicates collective movement by forcing group members to modulate their speed in order to maintain group cohesion. Our agent-based models demonstrate that to replicate the levels of cohesion we observe in wild animal groups, group members need to dynamically adjust their patterns of movement in response to their social context. Simultaneous tracking of the majority of a group of wild baboons using GPS and accelerometer data loggers provided an opportunity to assess how individuals modulate their fine-scale behavior in response to changes in their social environment, and thereby maintain the spatial cohesion of their group. Individuals have a characteristic stride frequency that is related to their body size, but they adjust this stride to match the pace of movement of their nearest neighbors. Furthermore, individuals deviate more from their characteristic stride frequency when group spread increases. Individuals also balanced their tendency to pause during group movement as a function of their spatial position within the group, waiting when they outstripped the group, and hustling to catch up when they fell behind. While all group members modulated their movement patterns in these ways, they did so to differing degrees. Compared to other members of their group, small baboons showed larger deviations from their characteristic stride frequency (Figure 3B). Consistent with previous work suggesting that changes in gait characteristics have important energetic consequences (Hoyt & Taylor 1981), smaller baboons also had higher VeDBA (i.e. a proxy for energetic expenditure) than their larger group-mates. Size-based differences in VeDBA were magnified as travel speed increased (Figure 2B). These results suggest that small individuals pay a disproportionate share of the energetic costs associated with maintaining group cohesion. Small individuals may incur additional costs if the effort required to keep up with their group-mates decreases their foraging efficiency — a likely outcome if small individuals are unable to pause to ‘forage on the go’. Because small individuals are more vulnerable to predators (Cowlishaw 1994), they may benefit more from the protection that group cohesion affords. Our results are thus consistent with the hypothesis that the costs of maintaining cohesion are largely borne by individuals that have the most to gain from group membership.

Baboons modulate their fine-scale movement decisions differently depending on their spatial position within their troop. We showed that individuals at the front of the group are more sensitive to group spread and pause often to let the rest of their group catch up, while individuals at the back of the group allow more separation from the group before increasing their move:pause ratios to catch up. This likely reflects context-dependent costs and benefits of different relative positions within the group. Differences in spatial positioning create variation in the ability of group members to influence group-level decisions (Couzin *et al*. 2005; Petit *et al*. 2009; Farine *et al*. 2017; Mclean *et al*. 2018), which suggests that being at the back of the group may compromise an individual’s opportunity to contribute to the group’s consensus decisions. However, the willingness of individuals at the back of the group to allow a larger separation from other group-members may reflect the benefits incurred by pausing often for small foraging bouts (and thus falling behind), and may be enabled by a perceived lower risk of predation associated with this position. Conversely, when individuals outstrip their group, their decision to slow down likely reflects a tradeoff between the opportunity costs of delaying arrival at their destination and the benefits of pausing to forage, as well as maintaining proximity to group-mates while in this particularly risky position within the group (Hamilton 1971; Krause 1994; Ioannou *et al*. 2015, 2019).

While the baboon troop as a whole travelled an average of 10.2 km per day, individual group members had significant variation in their daily travel distances: on the same day, some baboons travelled up to 1.1 km further than other members of their troop. Because all members of the troop followed the same general route, these individual differences in travel distance result from variation in individuals’ local, small-scale movements. In general, smaller individuals had longer daily travel distances, suggesting that body size may play a role in the sinuosity of an individual’s track. Inter-group and inter-population differences in baboon troop daily travel distances are well studied and can be attributed to a range of social and environmental factors including group size, food availability, and local interactions with other baboon troops and other species (Dunbar 1992; Pebsworth *et al*. 2012; Johnson *et al*. 2015; Slater *et al*. 2018). However, study of fine-scale variation in the daily travel distances of individuals in heterogeneous, socially cohesive groups is lacking. To our knowledge, there is no theoretical framework that explains why such differences arise. A more in-depth study of the effects of group members’ body size, age-sex class, social rank, and affiliative network on the fine-scale differences in their daily travel distances is needed to understand why some individuals travel farther, even along the same route.

The differences in VeDBA documented in this study suggest that the energetic consequences of collective movement vary among members of heterogeneous groups and that moving as part of a group is, in general, more energetically costly than moving alone. It is well established that the ecological cost of transport is highly variable across species, ranging from 0.19% to 28% of overall energy expenditure (Garland Jr. 1983; Husak & Lailvaux 2017), and that smaller animal species have higher energetic costs associated with locomotion (Taylor *et al*. 1982). While less is known about intraspecific relationships between body size and the energetic costs of locomotion, studies suggest that the same holds within species (Sockol *et al*. 2007; Pontzer *et al*. 2011; Sankey *et al*. 2019). This is consistent with our results showing that VeDBA was higher for smaller individuals and that the relationship between increasing speed and increasing VeDBA scaled with body size, with smaller individuals having relatively higher increases in VeDBA over increasing speeds. Variation in energy expenditure can result not only from variation in travel speed, but also from differences in individuals’ tendencies to move and pause (Kramer & McLaughlin 2001), as well as variation in the cost of turning while moving (Wilson *et al*. 2013). However, energy expenditure encompasses only a part of an individual’s energy balance. It is yet to be revealed how metabolic rates, energy intake, endurance and recovery dynamics change with body size (Nagy 2005; Birat *et al*. 2018), but all could potentially impact the cost of collective motion in heterogeneous groups. Tri-axial accelerometry provides a promising new method of quantifying many such inter-individual differences and may afford new opportunities for studying the costs of sociality in wild animals.

Heterogeneity in locomotor capacities between group members can have implications on the energetic demands of its members and may constrain group size and composition. First, larger groups must travel farther each day to meet their energetic needs (Clutton-Brock & Harvey 1977; Majolo *et al*. 2008), yet longer daily travel distances exacerbate inequalities in the energetic costs of locomotion. Increasing group size would thus lead to increasing disparities in the energetic expenditures of group members. The maximum daily travel potential of the smallest individuals could therefore contribute to limiting group size within a given species.

Second, the magnitude of the differences in locomotor capacities between the smallest and largest individuals of a given species may constrain the extent to which heterogeneous social groups can be cohesive. If the smallest and largest age-sex classes of individuals have such great disparities in locomotor energy expenditure that smaller individuals simply can’t keep up with, or travel as far as, larger individuals over the course of full days, this could cause a reduction in groups’ spatial cohesion, or even fission-fusion social dynamics (e.g., Pontzer & Wrangham 2006).

The compromises that individuals make to maintain group cohesion occur across many axes – including compromises related to dietary, safety, and social needs (Krause & Ruxton 2002; Markham & Gesquiere 2017; Jolles *et al*. 2020). A holistic view that considers the interactions between these axes of compromise is necessary to understand how individuals balance the costs and benefits of group-living. In baboons, group movement trajectories are steered by a process of shared decision-making among group members, suggesting that individuals may often make compromises in the timing and direction of movement in order to stay with their group (Strandburg-Peshkin *et al*. 2015). In this study, we show that individuals modulate their fine-scale locomotor behaviors relative to their social context and spatial position within the group during collective movement. All group members thus make locomotor compromises to maintain group cohesion, suggesting that the *pace* of collective movement is also driven by a shared decision-making process. Our findings stress the importance of considering the interaction between social dynamics and locomotor capacity in shaping the movement ecology of group-living species, and illustrate an approach for accomplishing this under socially and ecologically relevant field conditions.

## Acknowledgements

We thank Kenya National Science and Technology Council, Kenyan Wildlife Service and Mpala Research Centre for permission to conduct research. We are grateful to Alison Ashbury, Tanya Berger-Wolf, Damien Farine, Ariana Strandburg-Peshkin and Mark Grote for helpful comments and suggestions on earlier drafts. We thank M Wikelski, E Bermingham, D Rubenstein, M Kinnaird, D Carlino, and B Tripard for logistical support; R Kays, S Murray, M Mutinda, R Lessnau, S Alavi, J Nairobi, F Kuemmeth, W Heidrich, D Pappano, I Brugere, and J Li for field assistance. We acknowledge funding from the Max Planck Institute for Animal Behavior, the Smithsonian Tropical Research Institute and University of California. RH and MCC were supported by an NSF grant (IIS 1514174). MCC received additional support from Packard Foundation Fellowship (2016-65130), and the Alexander von Humboldt Foundation in the framework of the Alexander von Humboldt Professorship endowed by the Federal Ministry of Education and Research awarded to MCC. JCL was supported by an NSF Graduate Research Fellowship and a UC Davis Dean’s Distinguished Graduate Fellowship. Support was also provided by the Center for the Advanced Study of Collective Behavior at the University of Konstanz, DFG Centre of Excellence 2117 (ID: 422037984).

## Ethics Statement

All procedures were subject to ethical review and were carried out in accordance with the approved guidelines set out by the National Commission for Science, Technology and Innovation of the Republic of Kenya (NACOSTI/P/15/5727/4608). Baboon tracking was approved by the Smithsonian Tropical Research Institute (IACUC 2012.0601.2015).

## Video legends

Video 1. Observed and simulated groups moving in a single dimension. Individuals with longer leg length are represented in green, and shorter, in blue.

**Figure S1.**
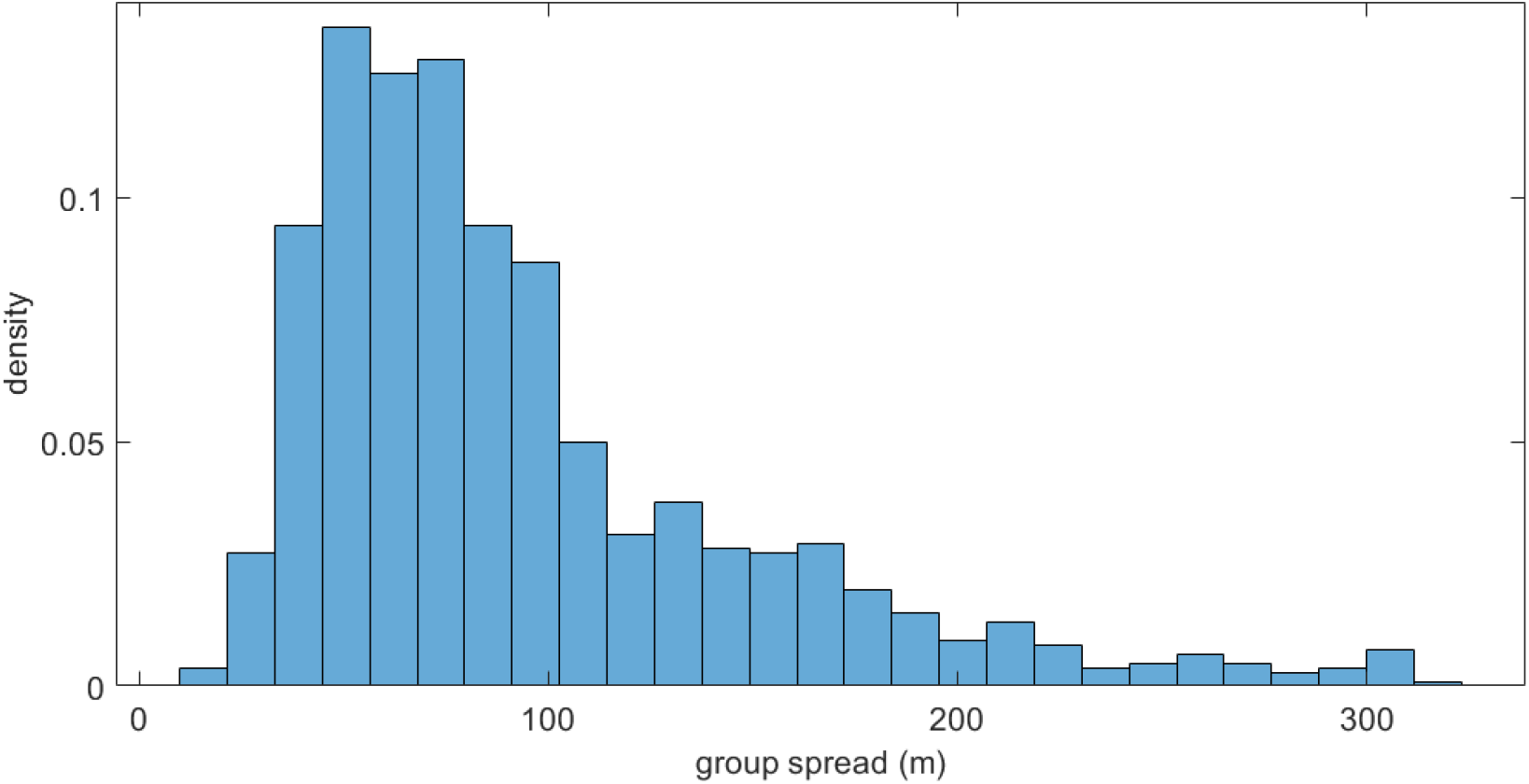
Group spread patterns. Group spread ranged between 35 and 320 meters.

